# Transient K^+^ current explains cross-species differences in the effects of myofibroblasts on myocytes

**DOI:** 10.1101/2020.11.02.365650

**Authors:** Fusheng Liu, Hou Wu, Xiaoyu Yang, Yuqin Dong, Guoyou Huang, Guy M. Genin, Tian Jian Lu, Feng Xu

## Abstract

Electrical and paracrine couplings between cardiomyocytes (CMs) and myofibroblasts (MFBs) affect both physiology and pathophysiology of cardiac tissues in a range of animal models, but relating these observations to humans is a challenge because effects vary greatly across species. To address this challenge, we developed a mathematical model for mechanoelectrical interactions between CM and MFB, considering both electrical and paracrine couplings between CMs and MFBs, with the aim of identifying the sources of cross-species variation and extrapolating animal models to predicted effects in humans. Our results revealed substantial differences across species in how these couplings modulate excitation-contraction coupling and Ca^2+^ transients of CMs. Both classes of couplings prolong action potential and increase APD in rat CMs, but shorten action potential and decrease APD in human CMs. Electrical coupling attenuates Ca^2+^ transients and active tension generation in human CMs, but has no significant effect on rat CMs. Paracrine coupling reduces Ca^2+^ transients and active tension in both human and rat CM. The results suggest that the variance of functional interactions between CM and MFB in cross-species may be explained by differences in the transient outward K^+^ currents associated with the KCND2 gene, and thus suggest potential therapeutic pathways for fibrotic cardiomyopathy.

## 1. Introduction

Cardiac fibrosis is a major source of morbidity and mortality arising from the interference of cardiac cardiomyocytes (CMs) with myofibroblasts (MFBs) and from excessive deposition of extracellular matrix by MFB (1, 2). Fibroblasts, quiescent in healthy cardiac tissue, can differentiate into MFB in the injured heart, particularly in the border zone of infarcts (3). MFBs can interfere with the contractile function of CMs through electrical coupling via gap junctions (*e.g*., Connexin 43, Cx43) (4) and through paracrine coupling (*i.e*., paracrine communication) via the secretion of paracrine factors (*e.g*., TGF-β1) (5). Understanding how MFBs modulate the mechanoelectrical behavior and calcium signaling of CMs is crucial to understanding the progression of cardiac fibrosis.

The evidence for these couplings between CMs and MFBs is now strong. Electrical coupling between CM-MFB by gap junctions has been established *in vitro* (6), but interleukin-1 β released by MFBs downregulates Cx43 (7). This had led to speculation that MFB-CM coupling does not occur in the native cardiac tissue, but the recent observations of Rubart *et al*. (8) confirm MFB-CM electrical coupling via Cx43 *in vivo*. Paracrine factors released by MFBs can change the volume fraction of T-tubule (9), the density of Na^+^ and K^+^ channels of ventricular myocytes (5), and the density of L-type Ca^2+^ channel (10) in atrial myocytes through regulating their mRNA expression. However, the ways that these electrical and paracrine factors work synergistically to affect cardiac function are not clear. Models for CM-MFB electrophysiological interactions are relatively advanced. MacCannell *et al*. (11) and Sachse *et al*. (12) have employed mathematical models to investigate the electrophysiological effects between CM-MFB through electrical coupling in human and adult rat. However, their results appear to contradict, where action potential of human CM coupled with MFBs tends to shorten (11) and that of rat CM coupled with MFBs tends to elongate (12).

This study aims to combine models of electrical and paracrine effects of MFBs to shed light on these differences, and to combine these with electrophysiological and contractile models of human and rat ventricular CM to assess the effects of MFBs on the calcium signaling and contractility of CM. Our results were validated by recent experimental findings. When the effects of electrical, paracrine, and synergistic couplings on mechanoelectrical function and Ca^2+^ transients of rat and human CM were compared, several potential sources for variance amongst species emerged.

## 2. Methods

### 2.1 Electrophysiological model of MFBs

MFBs have certain biophysical characteristics similar to CMs. Fast Na^+^ channel current (I_Na_), voltage-gated outward K^+^ channels current (I_K_), and mechano-gated channels current (I_MGC_) have been measured from mice and human MFB, where K^+^ channel proteins are expressed over >71% of all and regulate the main electrophysiological properties of MFB(13). Here our built a novel model to describe the electrophysiology of MFB. The model included six types of current. The model equation of In and the mechano-gated channels current (I_MGC_) were developed based on the newly experiments (14, 15). The Sachse *et al* (12) equations of inwardly rectifying K^+^ channel current (I_K1_) and voltage-gated outward K^+^ current (I_Kv_) were adopted to our model because it is closer to MFB K^+^ current relationship in experiment. The Na^+^-K^+^ pump current (I_NaK_) is necessary for the balance of intracellular Na^+^ influx and K^+^ effluxes in MFB. The I_NaK_ equation of MacCannell *et al* (11) was used in our model. Furthermore, a background current (I_b_) was added to responsible for the resting transmembrane potential, including background Na^+^ current (I_bNa_) and background K^+^ current (I_bK_). Ignoring the microscopic MFB structure, the cell membrane was model as a capacitance connected in parallel various channel currents and pumps. The electrophysiology of MFB is described by H-H equation,

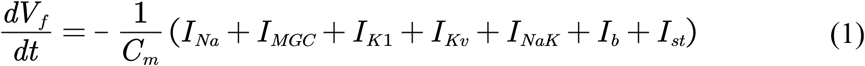

Where V_f_ is the transmembrane potential of a MFB, I_st_ is the stimulus current. C_f_ is membrane capacitance with the value 18 pF, according to the experiment measurement (9, 16).

### 2.2 Mechanoelectrical model of left ventricular CM

Pandit *et al* (17) and ten Tusscher *et al* (18) electrophysiological models of the adult rat and human left ventricular CM are reliable and widely recognized. They was built based on massive and widespread experimental data and widely used to research cardiac disease. Here we added the mechanical part of Mullins model (19) to Pandit and ten Tusscher model to construct a combined mechanoelectrical model of rat and human CM. Meanwhile, the kinetics equation of Ca^2+^ binding to troponin C from Mullins model was added to describe excitation-contraction coupling. Some parameters were modified based on the newly experimental measurements.

### 2.4 Coupling between CM and MFBs

Both electrical coupling mediated by Cx43 (**Fig. 1A**) and paracrine coupling mediated by TGF-β1 released by MFB (**Fig. 1B**) were considered. Electrical coupling regulates the direct electrophysiological interactions of the CM and MFB. The conductivity G_gap_ of Cx43 is close to linear function of contact lengths between CM and MFB (16). We assumed that the mean effective contact length is 5 μm, and then the G_gap_ value is 7 nS according to 1.4±0.6 nS/μm. The electrical coupling differential equation is governed by,

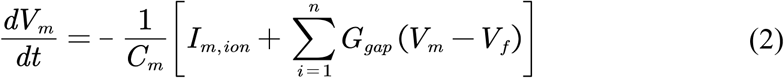

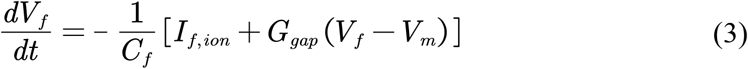

**Figure 1.**
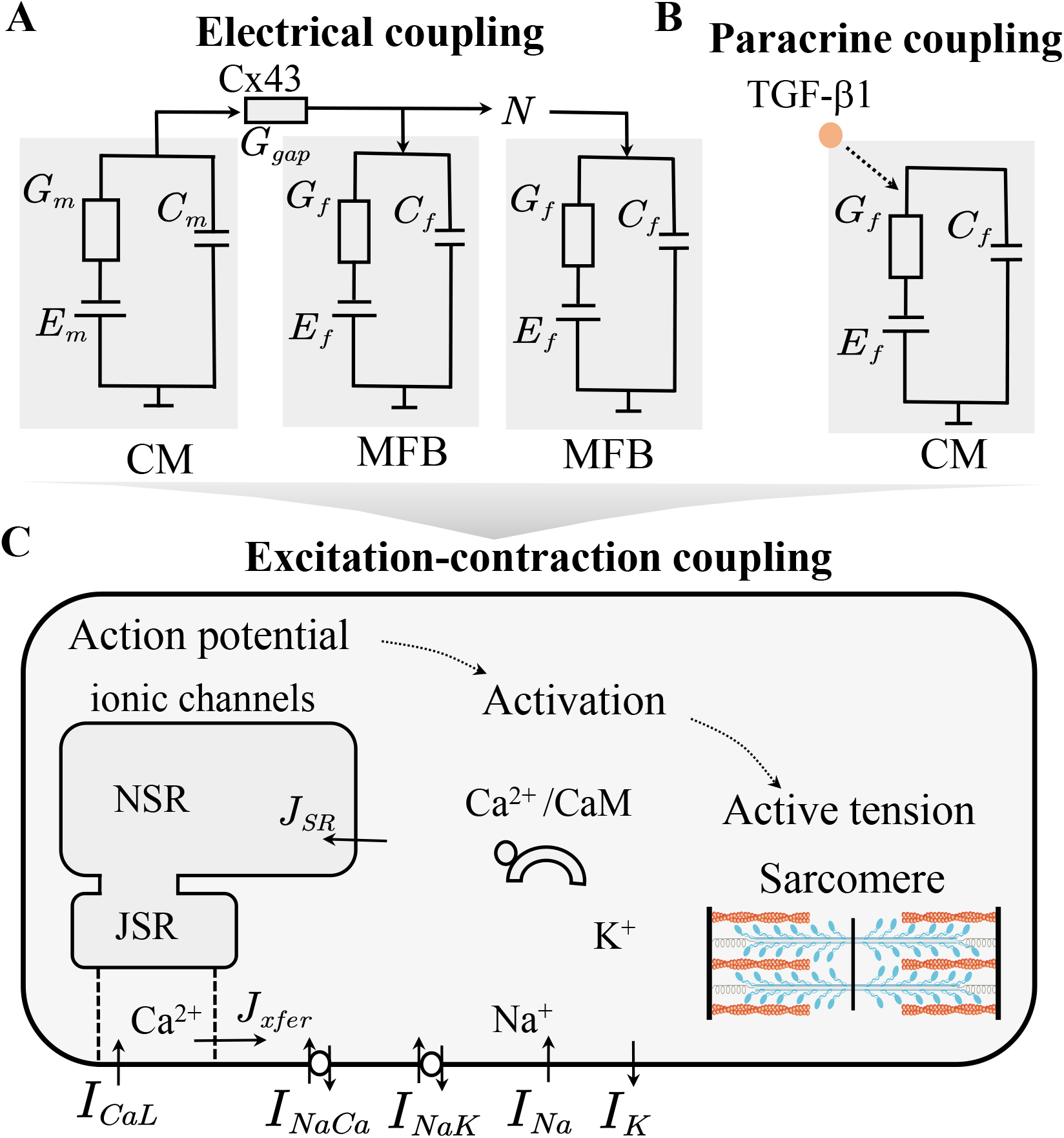
Functional interactions between MFB and CM by electrical and paracrine couplings. (A) Schematic of electrical coupling between CM and MFB that modulate the electrophysiology of CM as mediated by Cx43 connections. (B) Schematic of paracrine coupling between MFB and CM, as mediated by paracrine factors (*e.g*. TGF-β1). Paracrine factors, released by MFB, alter ion channels and T-T-tubules in CMs, and thereby modulate CM electrophysiology. (C) Electrical or paracrine coupling modulates electrophysiology and thereby affects Ca^2+^ transients and contraction of CMs.

Where V is the transmembrane potential of a CM (V_m_) or a MFB (V_f_), C is the membrane capacitance of CM or MFB (C_m_ or C_f_), I_ion_ is the membrane current of them, and n is the number of MFB coupled with CM.

Paracrine factor (*e.g*., TGF-β1) released by MFB regulates ion channel currents and T-tubule volume fraction and thereby indirectly affects action potential and calcium release in sarcoplasmic reticulum of CM. TGF-β1 was report to change ion channels protein expression and construction of CM (5, 9, 10). TGF-β1 reduces gene expression of Cav1.2 and L-type Ca^2+^ channel current of atrial CM, but not significantly alters in ventricular CM (10). Gene expression of SCN5A and Na^+^ channel current are increased, but Gene expression of KCNIP2, KCND2 and *I_to_* are decreased by TGF-β1in ventricular CM (5). TGF-β1 may also reduce T-tubule volume fraction of ventricular CM which cut down the Ca^2+^ release probability from sarcoplasmic reticulum (9). The changes were simulated by the above mechanoelectrical model of ventricular CM.

These electrophysiological interactions regulate active contraction and calcium signaling of CM through calcium induced calcium release (**Fig. 1C**). Based on these, we developed an electrophysiological model for MFB, which was then integrated into prior electrophysiological and contractile models of human and rat ventricular CM.

### 2.5 Model simulation

All the equations were solved by ode15s and forward Euler difference method in MATLAB. The forward Euler difference method is performed with the maximum time step of 0.005ms. (MathWorks, Version R2020a). Description and validation of our model formulation and parameter definitions were provided in the **Supplemental Material** (**Figs. S1-6**).

## 3. Results

### 3.1 Species-dependent functional interactions of CM and MFB via electrical coupling

To assess the effect of electrical coupling on functional interactions between CM-MFB in different species, a left epicardial ventricular CM of rat or human was coupled with zero, one, three or six MFBs at a cardiac frequency of 1 Hz (**Fig. 2**). The increased number of MFBs causes the accelerated repolarization, decreased repolarized plateaus, and attenuated action potential of human CM (**Fig. 2A**). In addition, increasing MFBs induces a significant decrease of Ca^2+^ transients (**Fig. 2B**) and active tension (**Fig. 2B-C**) in human CM. However, the increased number of MFB prolongs the action potential of rat CM (**Fig. 2D**). These electrophysiological results are in accord with reports from MacCannell *et al*. (11) and Sachse *et al*. (12). The number of MFBs has no significant effect on the Ca^2+^ transients and active tension of rat CM (**Fig. 2E-F**). This simulated Ca^2+^ transients of rat CM was verified by experimental observations from the co-culture of CMs and MFBs with TGF-β1 inhibited (**Fig. 2E**) (20).The trends of action potential of MFBs are consistent with the species of CM (**Fig. S7**). Most interestingly, we found that the action potential trend of CM coupled with MFBs in large mammals (*e.g*., rabbit, guinea pig) is similar to human and that of small mammals (*e.g*., mouse) is similar to rat (**Fig. S8**). We further compared the action potential duration (APD, 90% repolarization) restitution of human and rat CM coupled with three MFBs (**Fig. 2G**). We found that, at the same diastolic interval (DI), APD of human CM coupled with MFBs decreases but APD of rat CM coupled with MFBs increases.

**Figure 2.**
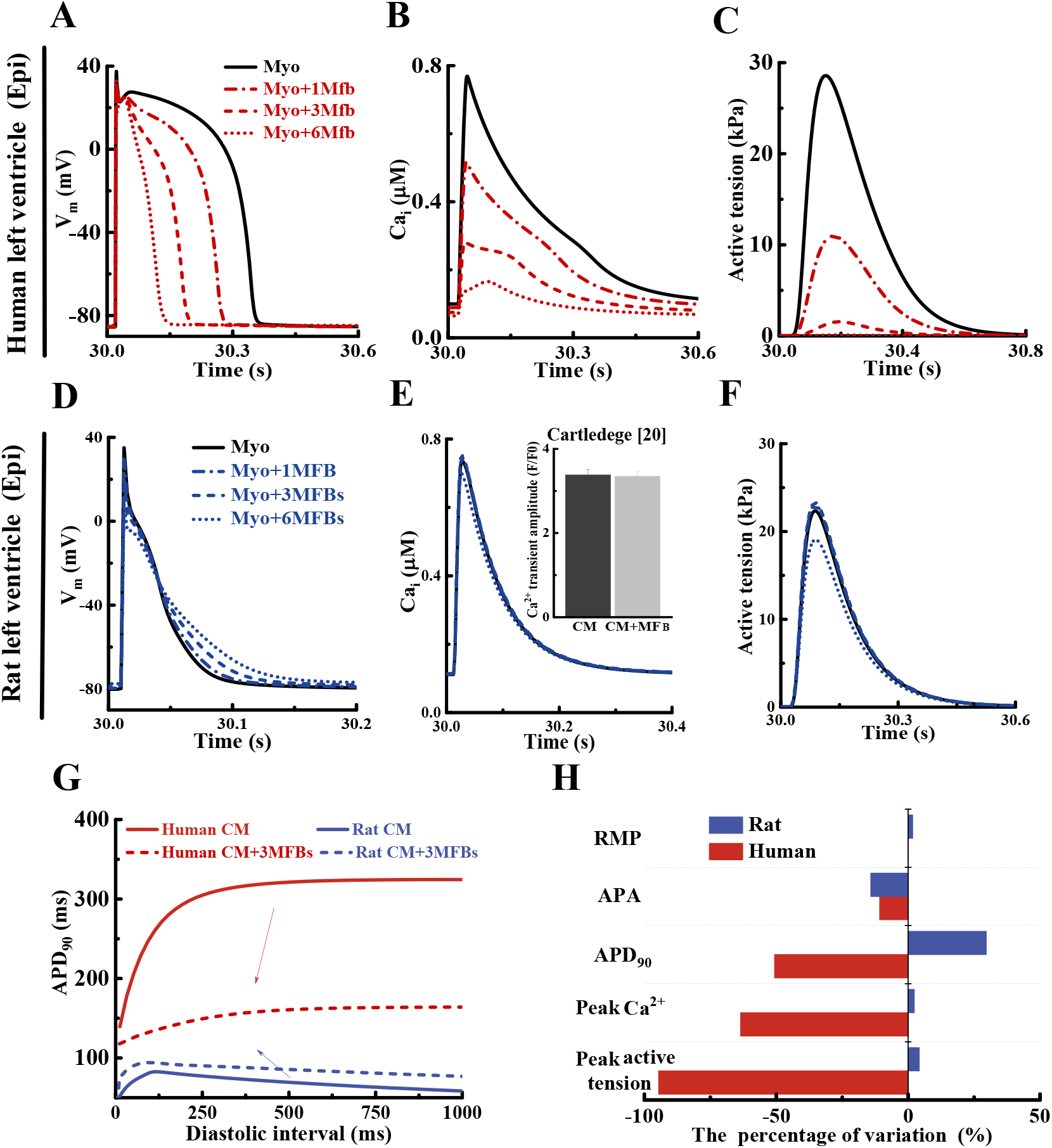
Electrical coupling with MFB modulates the function of human and rat CMs. Steady state action potentials V_m_ (A, D), Ca^2+^ transients (B, E), and active tensions (C, F) of human or rat CMs coupled with 0, 1, 3, or 6 MFBs. (G) APD_90_ restitution curves of a human or rat CM coupled with three MFBs, but with no interactions between CMs and MFBs, by the dynamic S1-S2 protocol with a second basic cycle. (H) Effect of electrical coupling on human and rat CMs coupled with three MFBs.

Figure 2H summarizes the functional variance of CM coupled with three MFBs compared to CM without interactions with MFB. For instance, resting membrane potential (RMP) of human CM and rat CM has barely changed. Action potential amplitude (APA) of them declines by 11% and 14%, respectively. Electrical coupling shortens the APD of human CM by 50%, but prolongs the APD of rat CM by 30%. Furthermore, peak Ca^2+^ transients and peak active tension of human CM decrease by 63% and 95%, respectively, but conversely those of rat CM improve by 2% and 4%, respectively. These results suggest that the mechanoelectrical interactions and calcium signaling between CM and MFB via electrical coupling reveal visible difference across species.

### 3.2 Species-dependent functional interactions of CM-MFB via paracrine coupling

To assess the effect paracrine factors released by MFB on CM function in different species, CM cultured with MFB conditional media was simulated (**Fig. 3**). TGF-β1 as a main paracrine factor was quantitatively measured in the culture media of ventricular CM and MFB by Cartledge *et al*. (20). The TGF-β1 concentration is about 30 times more in MFB than in CM (**Fig. 3A**). **Figure 3B-C** summarize the structural and functional effects of paracrine factor on ventricular CM from two recent experiments (5, 9), where fast Na^+^ current I_Na_ at −40 mV is increased by 40%, transient outward K^+^ current I_to_ at 60 mV and T-tubule volume fraction are decreased by 55% and 30%, respectively. These changes cause increased APD of human CM and reduced APD of rat CM, which agree with the experiments in large and small mammals (5, 9) (**Fig. 3D**). Then, we summarize the functional variance of CM induced by paracrine factor compared to CM without paracrine factor (**Fig. 3E**). There is no significant change of RMP for both human CM and rat CM. APA of human CM and rat CM improves by 5% and 10%, respectively. APD of rat CM improves by 47%, but APD of human CM decreases by 3%. Furthermore, peak Ca^2+^ transient of human CM and rat CM decrease by 28% and 13%, respectively, and maximal active tension decreases by 54% and 16%, respectively. These results suggest that the mechanoelectrical interactions and calcium signaling between CM and MFB via paracrine coupling exhibit obvious difference across species. We next simulated the effect of different TGF-β1concentrations on the action potential of rat CM. Action potentials is prolonged with increasing TGF-β1concentrations (**Fig. 3F**) which coincides with the experimental result from the literature (5).

**Figure 3.**
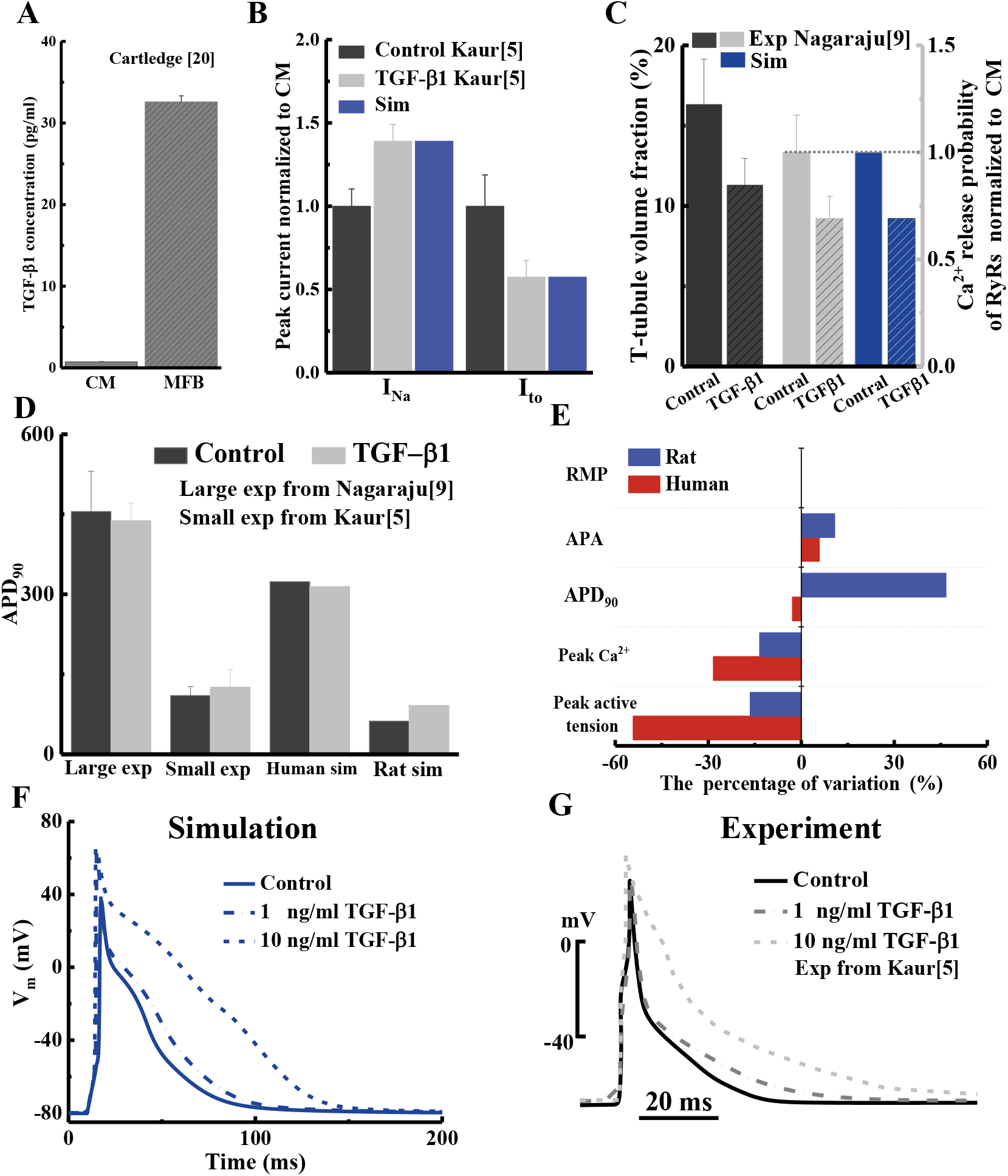
Paracrine coupling with MFBs modulates function of human and rat CMs. (A) Concentration of TGF-β1 in rat CM culture medium and MFB culture medium. (B-C) TGF-β1 increases peak fast Na^+^ current I_Na_ and reduces peak transient outward K^+^ current I_to_ and the T-tubule volume fraction. (D) Comparison of simulated APD_90_ in human and rat CMs and experimental measurement of APD_90_ modulated by TGF-β1in large and small mammal CM. (E) The effect of TGF-β1 on human and rat CMs. (F) Simulated and (G) experimental rat action potentials at different TGF-β1 concentration.

### 3.3 Synergistic effects of electrical and paracrine couplings

Synergistic electrical and paracrine couplings between CM and CFB were modeled by adding TGF-β1 from MFB culture media, and studying human CM and rat CM coupled with three MFBs (**Fig. 4**). Synergistic coupling significantly shortens action potential of human CM but prolongs that of rat CM (**Fig. 4A**). APD_90_ of human CM decreases by 51% but that of rat increases by 47%, as verified by co-culture experiments in large mammals (pig) (9) and small mammals (mouse) (21) (**Fig. 4B**). The Ca^2+^ transient and active tension are attenuated in both human CM and rat CM by synergistic coupling (**Fig. 4C and 4E**), with greater attenuation in human CM. The Ca^2+^ transient amplitude of rat CM agrees well with rat co-culture experiments (20) (**Fig. 4D**).

**Figure 4.**
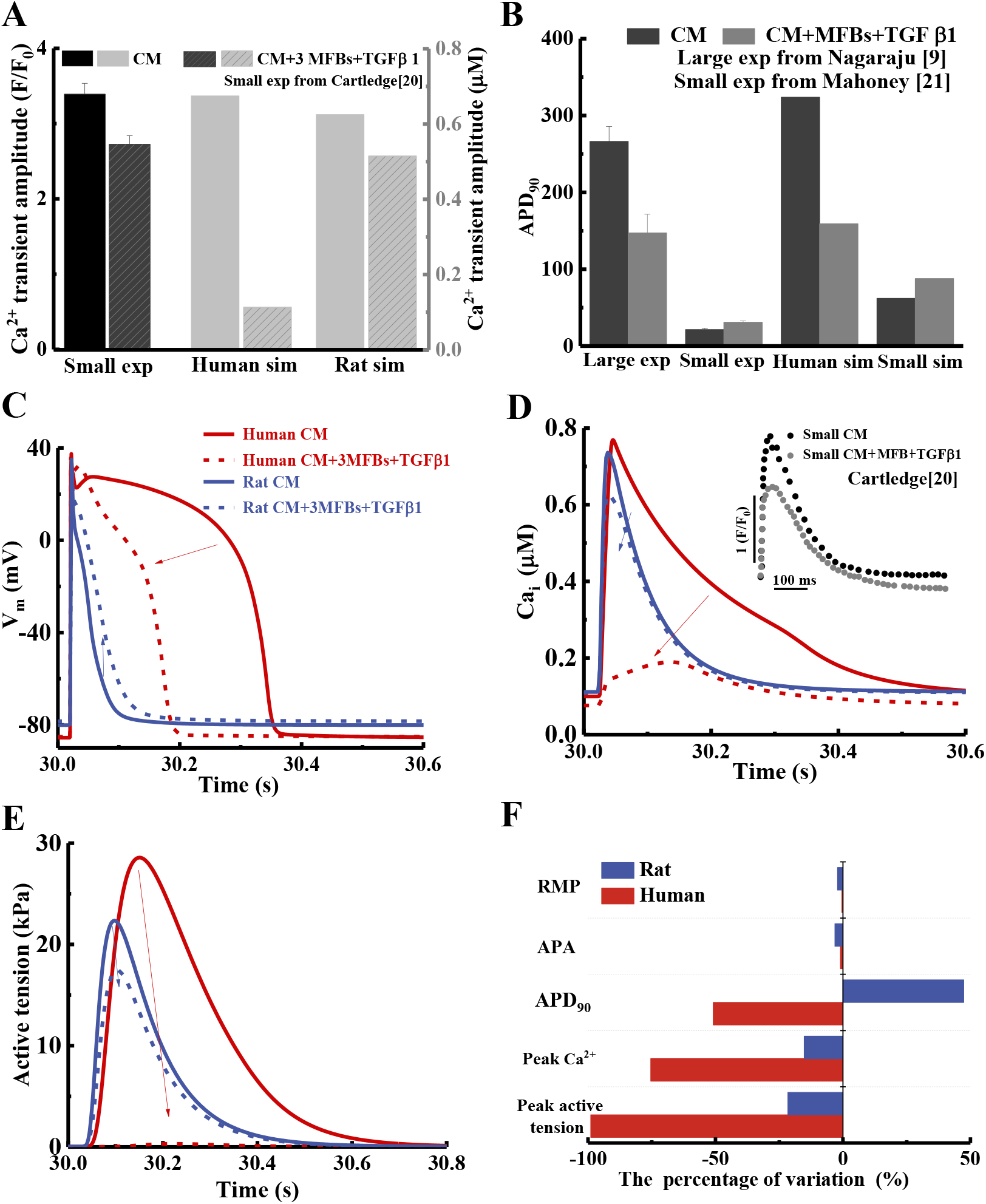
Synergistic coupling of MFB modulates the function of human and rat CM. Action potentials (A), Ca^2+^ transients (C) and active tensions (E) of CM and CM synergistically coupled with MFBs in human and rat. (B) Comparison of simulated and experimental measurement of APD_90_ in large mammalian CM (*e.g*., human, pig) with that in rat CM with and without interactions with MFB. (D) Comparison of simulated and small mammalian experimental Ca^2+^ transient amplitude of CM and CM synergistically coupled MFB between human and rat. (F) Effect of synergistic coupling on human and rat CM.

The effects of synergistic coupling (**Fig. 4E**) show no significant effects on RMP and APA in human CM and rat CM when compared to CM without interactions with MFB. APD_90_ increases by 47% in rat CM but reduces by 51% in human CM. Peak Ca^2+^ transient in human CM and rat CM decrease by 75% and 15%, respectively, and maximal active tension decreases by 99% and 21%, respectively. Compared to **Figures 2H, 3E**, and **4E**, we found the effects of synergistic couplings on CM are more significant than only electrical or paracrine coupling. This suggests that electrical and paracrine couplings between CM and MFB are collaborative processes. In addition, these results indicate that the functional interactions between CM and MFB by synergistic couplings also shows cross-species differences.

### 3.4 The *I_to_* current explains different CM-MFB interactions in human and rat

Human CM and rat CM behave differently in response to combined electrical and paracrine couplings, as reported by literature and our simulations. To explore the sources of the differences, we analyzed distinctions between human and rat CM ion channels (**Fig. 5A**).

**Figure 5.**
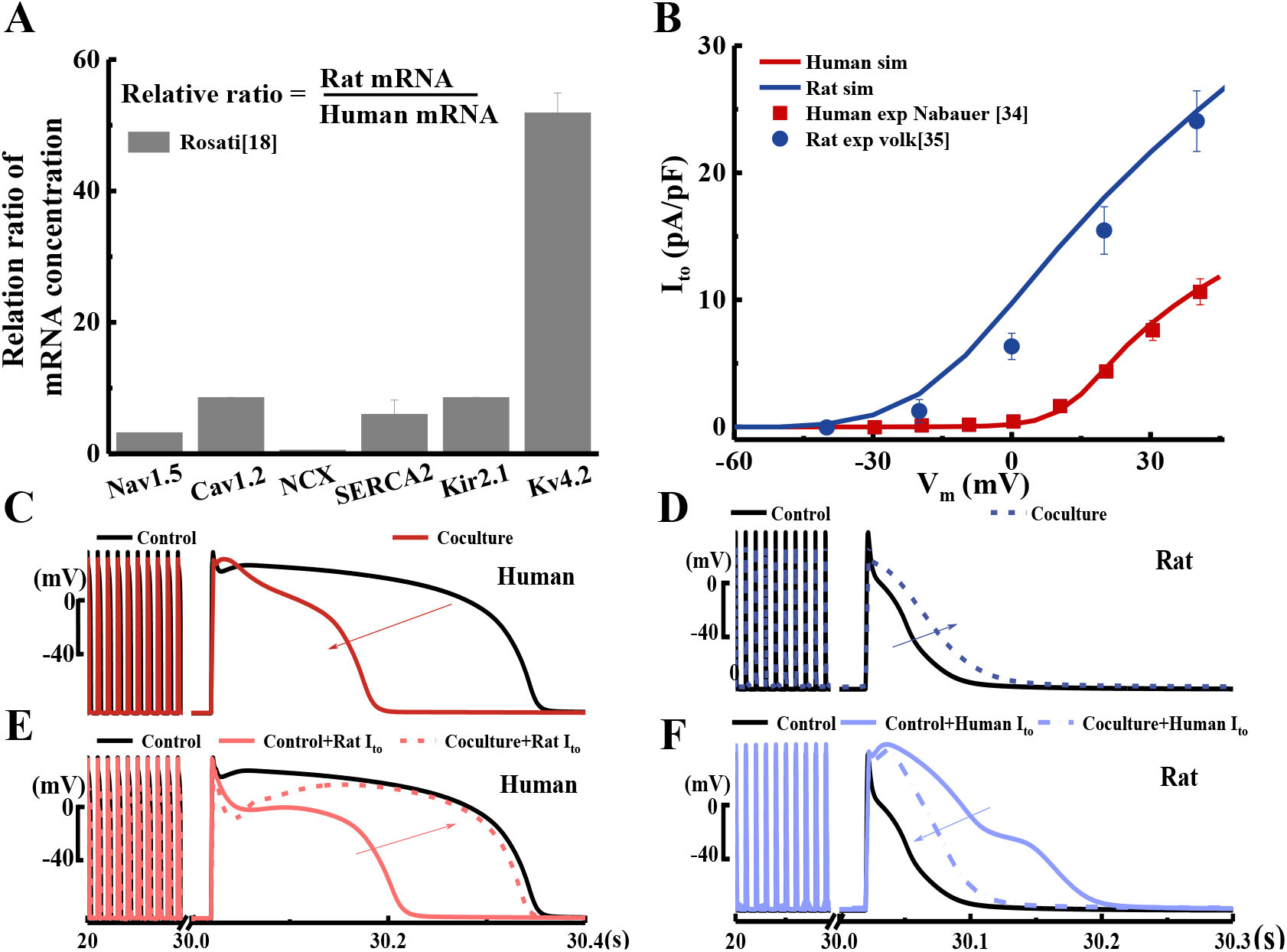
Gene expression for I_to_ modulates differences between the interactions of MFBs with human and rat CMs. (A) Relative ratio of main ion channel mRNA concentrations between human and rat CM. (B) Simulated and experimental I_to_ in human and rat left epicardial ventricular CMs. Synergistic coupling shortens the action potential of human CM (C) and prolongs that of rat CM (D). The action potential of human CM is prolonged (E) and that of rat CM is shortened (F) via the synergistic coupling after swapping models for human and rat I_to_ between human and rat CM.

We first explored genes associated with specific ion channels in human CM and rat CM respectively. In human CM, the KCNH2 and KCNQ1 genes produce the large slow delay rectifier K^+^ channel current (I_Kr_) and fast delay rectifier K^+^ channel current (I_Ks_); in rat CM, the HCN2,4 and KCNA5 genes produce the small hyperpolarization-activated inward K^+^ channel current (I_f_) and steady-state outward K^+^ channel current (I_ss_) (22). To assess whether these are the sources of the cross-species differences, we exchanged the two types of currents between human CM and rat CM and found that they do not change the trend of action potential: the action potential of human CM is still shortened and the action potential of rat CM is prolonged when CM is coupled with MFB (**Fig. S9**). This suggests that such specific ion channel currents across species (*i.e*., I_Kr_ and I_Ks_ of human CM and I_f_ and I_ss_ of rat CM) are not the key factors that contribute to species-dependent functional interactions of CM-MFB.

We next explored KCND2 gene (Kv4.2) that is associated with the transient outward K^+^ channel current (I_to_) expressed in both human CM and rat CM exhibit significant differences between human CM and rat CM. **Figure 5A** shows the relative ratio of mRNA concentration in the major channels between human CM and rat CM. KCND2 mRNA concentration is about fifty times higher in rat CM than in human CM (23), and the peak I_to_ current density is about 2.5 times higher in rat CM than in human CM (**Fig. 5B**). The reversal potential of human and rat I_to_ are 0 mV and −40 mV, respectively. We therefore next explored how I_to_ relates to the observed cross-species differences predicted for the synergistic electrical and paracrine effects of MFB. These factors explained the observed cross-species differences. Whereas synergistic electrical and paracrine effects of MFB shortened the action potential of human CM and prolonged action potential of rat CM (**Fig. 5C-D**), running the models with I_to_ exchanged between human and rat CM reversed this effect, with synergistic coupling prolonging action potential in human CM and shortening it in rat CM (**Fig. 5E-F**). The results suggest that the transient outward K^+^ channel current associated with the KCND2 gene may be a key factor underlying differences between human CM and rat CM responses to the synergistic electrical and paracrine effects of MFB.

## 4. Discussion and conclusions

Our model of how MFB affect human CM and rat CM excitation-contraction coupling suggests synergistic effects of electrical and paracrine couplings. The model, verified by experiments from the literature (5, 8, 20, 21) and by previous simulations (11, 12), suggest factors underlying differences amongst species in myocardial fibrosis (**Fig. 6**). The electrophysiology of CM controls Ca^2+^ transients and active tension of through excitation-contraction coupling, which contribute to electrical and paracrine couplings that affect the action potential, APA, APD and Ca^2+^ transient and active tension of CM. CM-MFB coupling tends to prolong action potential in rat CM but shorten action potential in human CM. Attenuation of Ca^2+^ transients and active tension in human CM is more obvious than in rat CM. Key factors underlying these differences are the transient outward K^+^ channel current associated with the KCND2 gene and the associated cross-species differences in I_to_.

**Figure 6.**
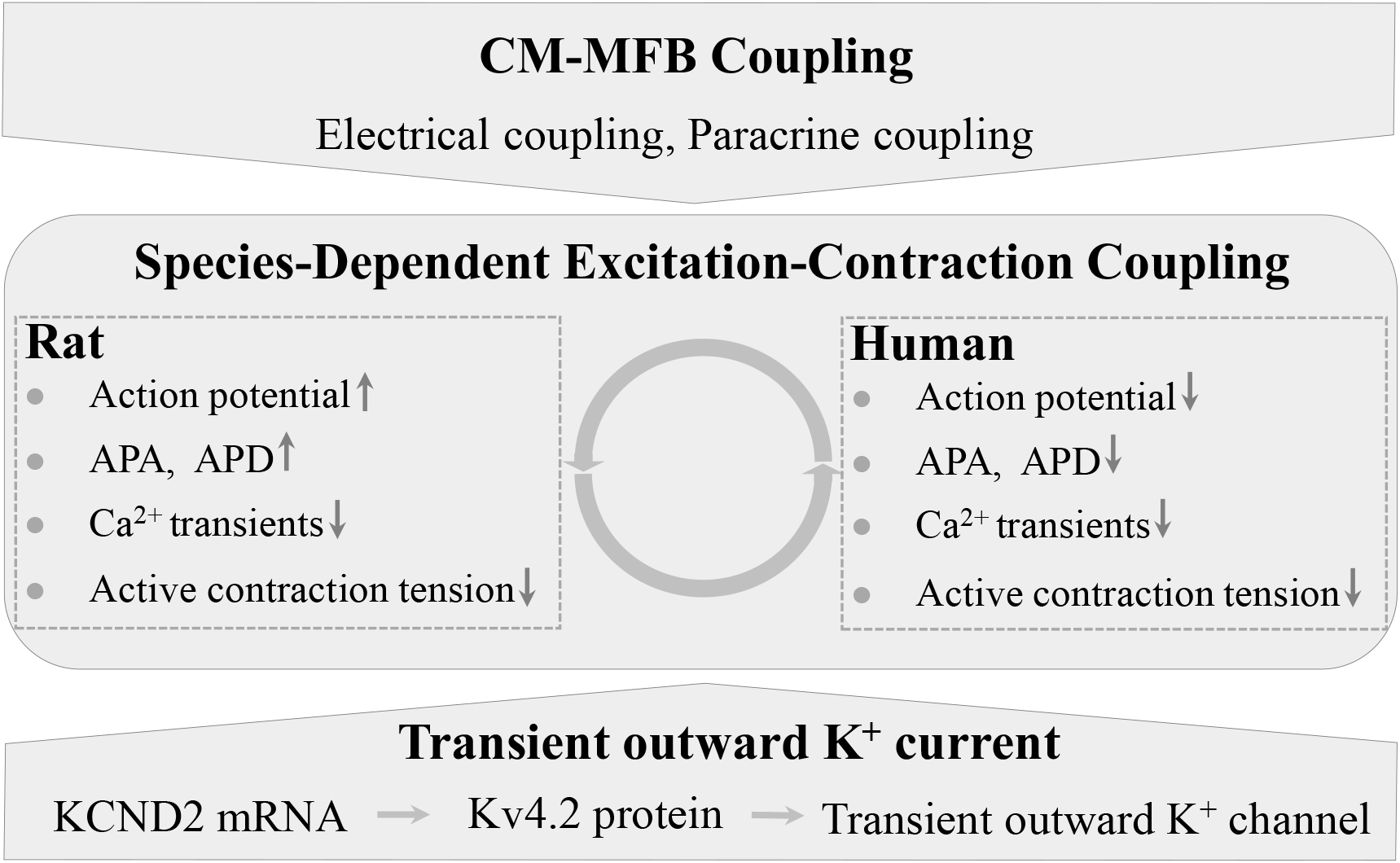
Species-dependent interactions between CM and MFB. Electrical and paracrine couplings regulate electrophysiological interaction between CMs and MFBs in myocardial fibrosis. Cardiac electrophysiology controls cardiac tension through excitation-contraction coupling. I_to_ associated with the KCND2 gene affects cross-species differences in interactions between CM and MFB.

### 4.1 The coupling mechanism between CM and MFB

Although others have assessed electrical coupling of MFB on electrophysiology and conduction of CM (11, 12, 24, 25), and reports of paracrine effects of MFB regulating ion channel expression and T-tubule of CM are widespread (5, 9, 10), this is the first study of which we are aware that simulates both factors and assesses their synergistic effects. Synergistic coupling between MFB and CM could alter the electrophysiology, Ca^2+^ transient and active tension of CM. Individual effects of electrical and paracrine signaling are similar, with the sole exception being that electrical coupling tends to weaken APA while paracrine coupling tends to enhance it because of increasing ?Na in CM caused by paracrine factor. Our simulations verify the observation that electrical coupling contributes to electrophysiological interactions in fibrotic myocardial scars (21). In addition, mechanical coupling, a underlying coupling mechanism between CM and MFB, may also affect electrophysiological conduction of CM (26) and contractility of MFB (27). The model therefore captures many results from the literature, and suggests additional experiments that can further dissect CM-MFB coupling mechanisms.

### 4.2 Species-dependent CM-MFB interactions

Action potentials, ionic currents and their mRNA expression vary across mammalian left ventricles (22, 23, 28-30), but it remains elusive what these differences mean physiologically. In this paper, we discovered that the action potential, Ca^2+^ transient and active tension in mammalian left ventricles exhibit changes in the presence of MFB that vary across species. CM-MFB coupling tends to prolong action potential in small mammals (*e.g*., murine) and shorten action potential in large mammals (*e.g*., rabbit, pig and human). Our results suggest that differences in transient outward K^+^ channel current may underlie these effects, with mRNA concentration of the underlying KCND2 about 50 times larger in small mammals than in large mammals (23). The peak I_to_ density of CM in small mammals is about 2.5 times of large mammals and the reverse voltage of I_to_ in large and small mammals are 0 mV and −40 mV, respectively (31). I_to_ modulates the early repolarization of action potential, which strongly impacts the time course and magnitude of Ca^2+^ transients and thereby affects cardiac contractile tension (32). Our results provide a mechanism for understanding how murine models of cardiac fibrosis relate to humans.

### 4.3 Limitations

Our computational work has a limitation that bears mention. Our electrophysiological model of MFB was developed on the basis of different species because of the limited experimental measurement of MFB. Our model of MFB includes six types of ion channels. A review by Feng *et al*. (33) showed the presence of more than nine types ion channels in MFB, particularly calcium associated channels. Calcium associated channels are key to excitation–contraction feedback, and quantitative experimental recordings of MFB ion channels remain a pressing need. Although experiments from the literature indirectly or directly verify our simulations, experimental data for electrical coupling between MFB and CM are limited, and we had to verify this part work by comparing it to previous simulations (11, 12).

However, the main findings of the study motivate new experimentation that may be able to quantify these CM-MFB interactions definitively. Electrical and paracrine couplings between CM and MFB are collaborative processes, and may affect electrophysiology, Ca^2+^ transients and active tension in ways that are species-dependent. The transient outward K^+^ current associated with the KCND2 gene and its effect on I_to_ are one key factor for cross-species differences, and may be a potential therapeutic target for myocardial fibrosis.

## Supporting material

Supplemental materials can be found online.

## Author contribution

F.L., G.G., T.J.L. and F X. conceived and designed the research project. F.L. H.W., Y.D and Y.Y. developed model methods and analysis, numerical coding and simulations. F.L., G. G. and F. X. written and revised the article. All authors reviewed the results and approved the final version of the manuscript.

## Acknowledgements

This research was supported by the National Natural Science Foundation of China (12032010, 11842015, 11972280, and 11872298).

## Supplemental Materials

### Model Description of Myofibroblasts (MFB)

#### 1. Fast Sodium Current Model of MFB

The newly measured experimental data of fast Na^+^ current of MFB was carried out by Chatelier *et al* at ~22°C (1). We used the traditional three gates formulation of I_Na_ to describe Na^+^ current, where there are an activation gate (*m*), two fast and slow inactivation gate (*h* and *j*). The steady-state curve for activation and inactivation, and the inactivation time constants were fitted to Chatelier et al (1) (**Fig. S1A-B)**. The activation time constants were from Luo and Rudy (**Fig. S1C**) (2). The simulation peak Na^+^ current were acquired with the same experimental test from a 500 ms holding potential of −120 mV to potentials ranging from −100 mV to 40 mV for 50 ms during [Na^+^]_o_ = 150mM (**Fig. S1D**). The maximal conductance of I_Na_ is 0.42 nS/pF and the R-squared coefficients is 0.99.

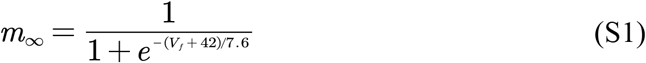

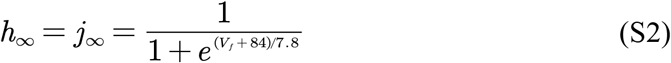

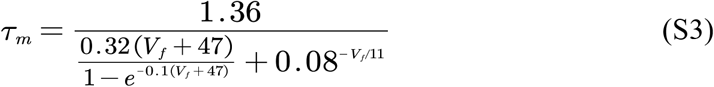

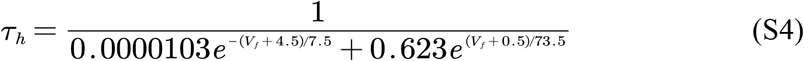

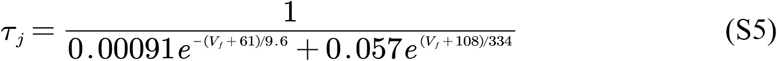

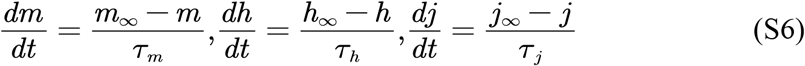

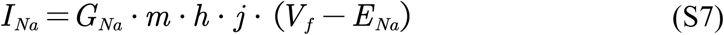

**Figure S1.**
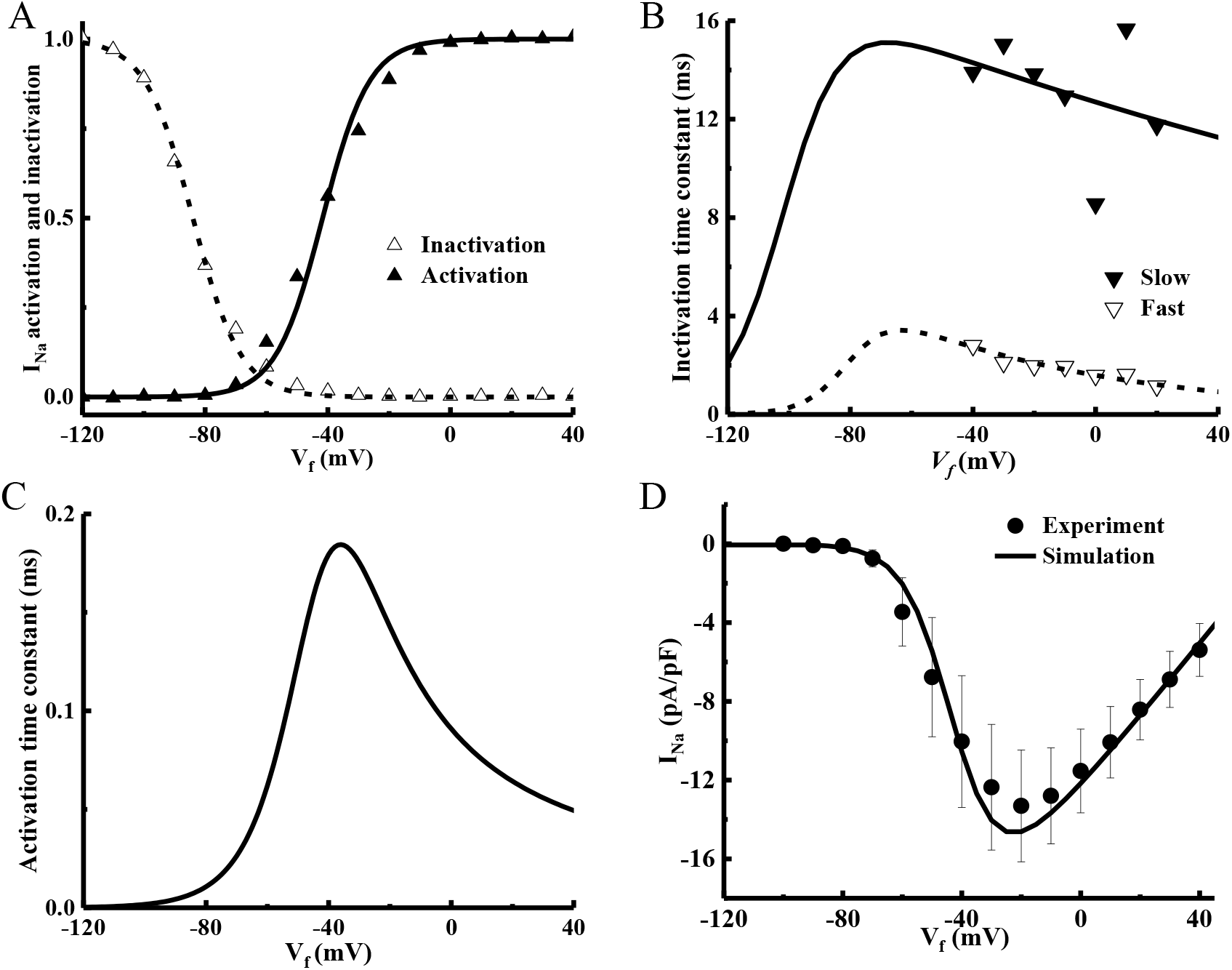
The validation of I_Na_ Model. (A)Steady-state curve for activation and inactivation. (B) Fast and slow inactivation time constants. (C) Activation time constants. (D) The simulated I-V curve of peak Na^+^ currents curve. Experimental data are from Chatelier *et al*(1).

#### 2. Mechano-gated channels current Model of MFB

The mechano-gated channels current I_MGC_ of MFB was recorded by Kamkin *et al* (3). There are two types of mechano-gated channels, a big I_MGC1_ and a small I_MGC2_. Their experiment implied that *I_MGC_* of MFB is activated by compressive deformation, but not stretches. The maximum compressive deformation causing completely opening of I_MGC_ is about 4μm. We used compressed length to describe the opening probability of I_MGC_ (**Fig. S2A)** because of the irregular shape of MFB. The opening probabilities of two types of mechano-gated channels are almost the same (3). The voltage-current relationship with the 3μm compressed length are shown in **Figure S2B**. The two maximal conductance of I_MGC_ are 0.012 nS/pF, 0.006nS/pF and the R-squared coefficients is 0.99.

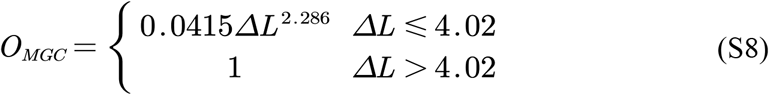

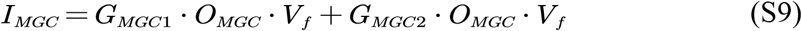

**Figure S2.**
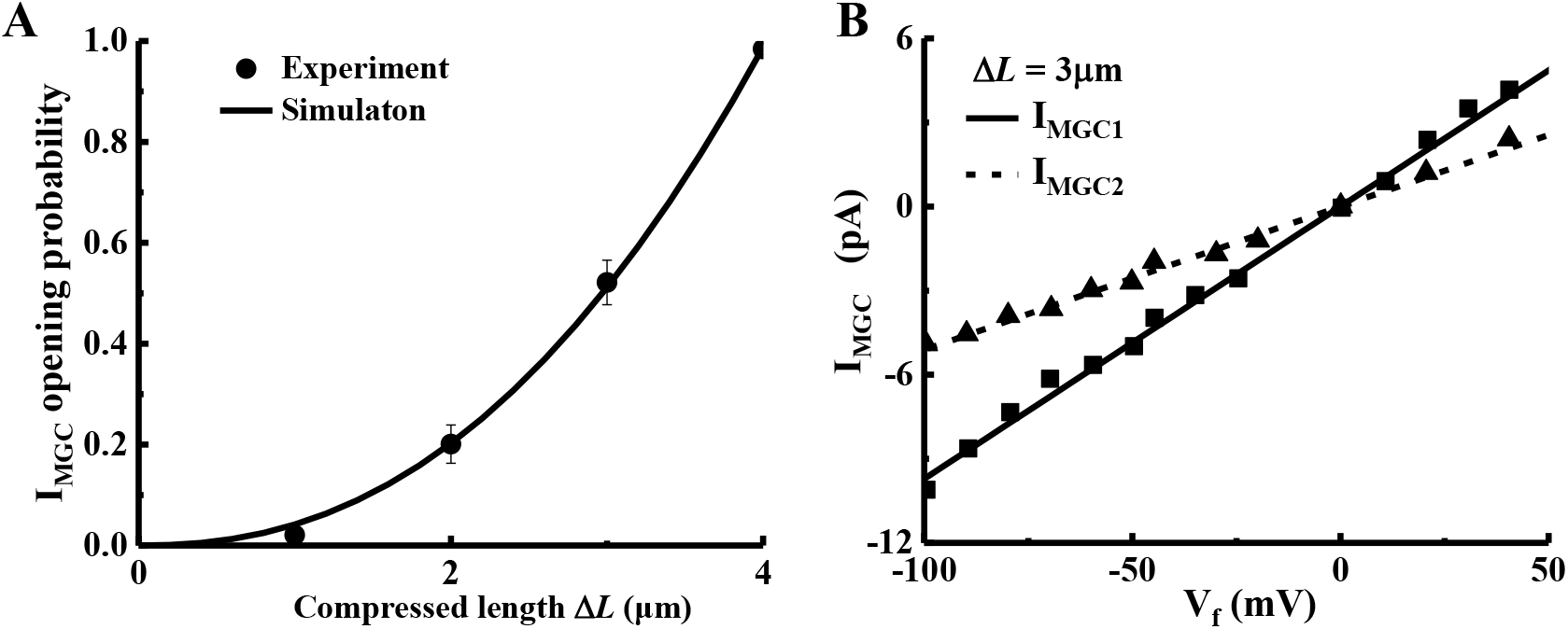
The validation of I_MGC_ Model. (A)The relationship of compressed length and opening probability of I_MGC_. (B) The voltage-current relationship with the different compressed length. Experimental data from Kamkin *et al*(3)

#### 3. Other channels current Model of MFB

The inwardly rectifying K^+^ channel current (I_K1_)and the voltage-gated outward K^+^ current (I_Kv_) of The Sachse *et al* (4) model were adopted to our model because it is closer to MFB K^+^ current relationship in experiment. The Na^+^-K^+^ pump current (I_NaK_) is necessary for the balance of intracellular Na^+^ influx and K^+^ effluxes in MFB. The I_NaK_ equation of MacCannell *et al* (5) was used in our model. Furthermore, a background current (I_b_)was added to responsible for the resting transmembrane potential, including background Na^+^ current (I_bNa_) and background K^+^ current (I_bK_). The two maximal conductance of IbNa and I_bK_ are 0.004 nS/pF and 0.005 nS/pF, respectively.

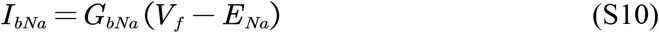

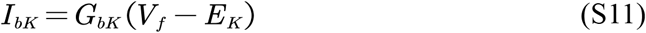

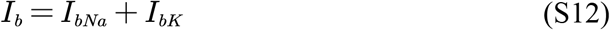

#### 4. Electrophysiological model validation for MFB

We built a novel MFB electrophysiological model based on Sachse *et al*(4) and MacCannell *et al*(5) opinion for MFB and newly experimental recordings (**Fig. S3**). The simulated action potential is in agreement with optical action potential recording (**Fig. S4A-B**). Simulated resting membrane potential (RMP) and action potential duration (APD_70_, 70% repolarization) are close to experimental measurements (**Fig. S3C**) (6, 7). **Figure. S3D** shows the comparison between the simulated and measured membrane voltage and outward K^+^ current curves. The reversal voltage improve from −40mV to −10mV at 10 and 100 mM extracellular K^+^ concentration. The simulation result of whole cell I-V curve are in accord with the Shibukawa *et al(6)* experiment at 22± 1°C with [K^+^]_o_ = 10mM and [K^+^]_o_ = 100mM (**Fig. S3D)**. The simulation results from our model have great agreements with experimental measurement.

**Figure S3.**
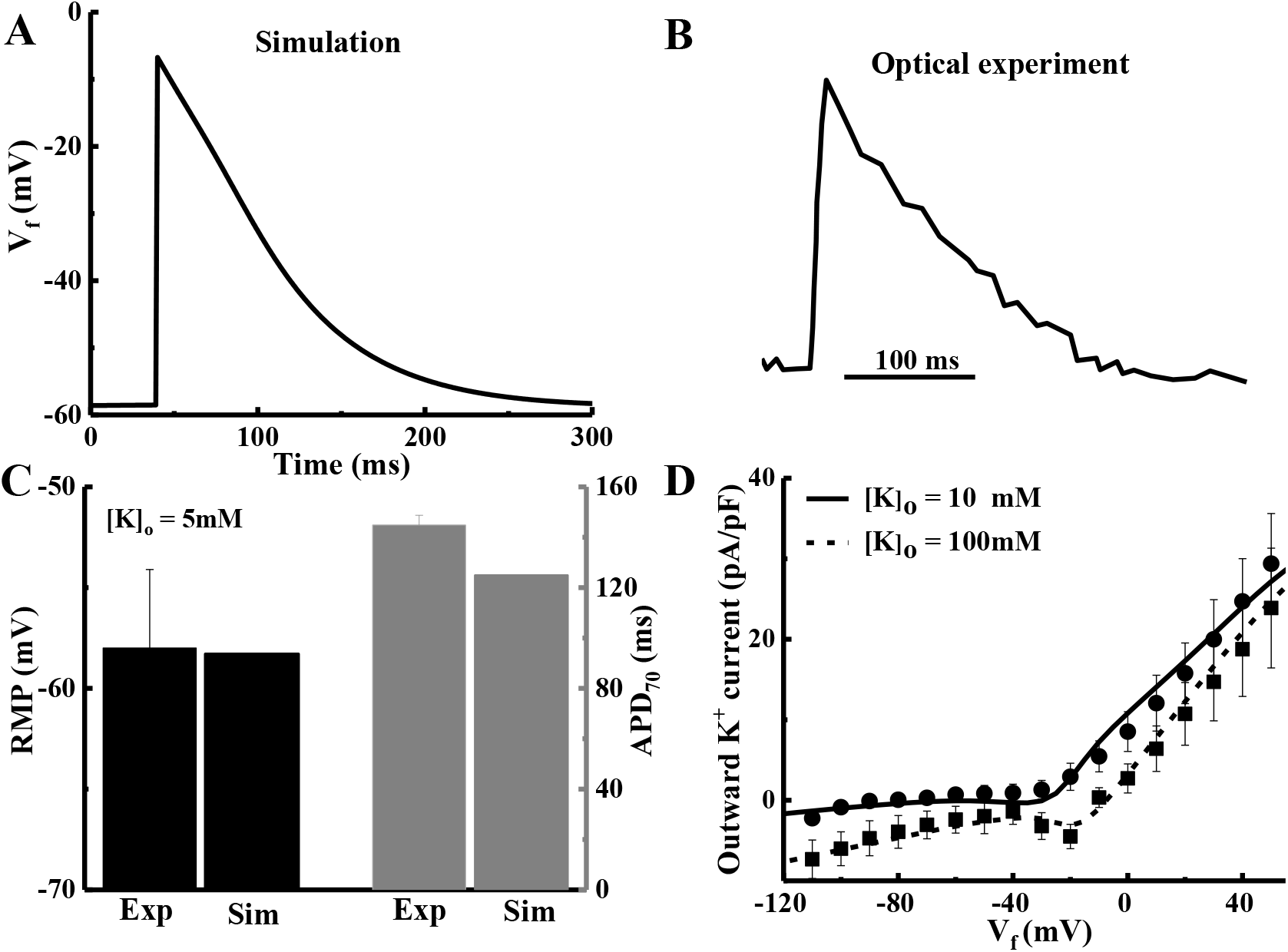
Validation of MFB model. (A) Simulated action potential of MFB under an externally applied stimulus −6 pA/pF for 5 ms. (B) Optical experimental action potential recording of MFB. (C) Comparison of simulated (line) and experimental (circle) I/V relationship for an extracellular K^+^ concentrations 10 and 100 mM. (D) Resting membrane potential (RMP) and APD_70_ from experiment and simulation. Simulated and experimental temperature is 22±1°C.

**Table S1.** Parameters of the MFB model.

| Parameter | Definition | Value |
| --- | --- | --- |
| T | Temperature | 295K |
| Cm | Cell capacitance of MFB | 18 pF |
| $[K^+]_i$ | Intracellular initial $K^+$ concentration | 140 mM |
| $[K^+]_o$ | Extracellular $K^+$ concentration | 5.4 mM |
| $[Na^+]_i$ | Intracellular initial $Na^+$ concentration | 10 mM |
| $[Na^+]_o$ | Extracellular $Na^+$ concentration | 140 mM |
| $G_{Na}$ | Maximal conductance of $I_{Na}$ | 0.42 nS/pF |
| $G_{Kv}$ | Maximal conductance of $I_{Kv}$ | $0.000354 \text{ cm}^3 \cdot \mu\text{F} \cdot \text{s}^{-1}$ |
| $G_{K1}$ | Maximal conductance of $I_{K1}$ | 0.178 nS/pF |
| $G_{MGC1}$ | Maximal conductance of $I_{MGC1}$ | 0.012 nS/pF |
| $G_{MGC2}$ | Maximal conductance of $I_{MGC2}$ | 0.006 nS/pF |
| $G_{bNa}$ | Maximal conductance of $I_{bNa}$ | 0.004 nS/pF |
| $G_{bK}$ | Maximal conductance of $I_{bK}$ | 0.005 nS/pF |
| $I_{NaKmax}$ | Maximal current generated of $I_{NaK}$ | 1.75 pA/pF |

#### 5. Comparison our MFB model and the prior models

Our model includes six types ionic channel to simulate further the electrophysiological behavior of MFB (**Table S1**). The capacitance of MFB (18 pF) and the conductivity between CM and MFB (14 nS) are bigger than Sachse *et al*(4) and MacCannell *et al*(5). We applied the same external electrical stimulus to our model with −6pA/pF for 5ms. The simulated action potential amplitude (APA) of MFB is far below that of CM under the same stimulus, which indicate MFB is weaker excitability than CM. The resting membrane potential (RMP) of our model is 58.6 mV, and the experimental RMP is 58.0±3.9 mV in [K^+^]_o_ = 5mM. Under an externally applied stimulus current −50pA/pF for 1ms, the action potential morphology of our model is similar to the Sachse model, but have not obvious plateau (**Fig. S4)**. Our simulated action potential have not obvious plateau and hook relative to MacCannell and Sachse model in the repolarization process.

**Figure S4.**
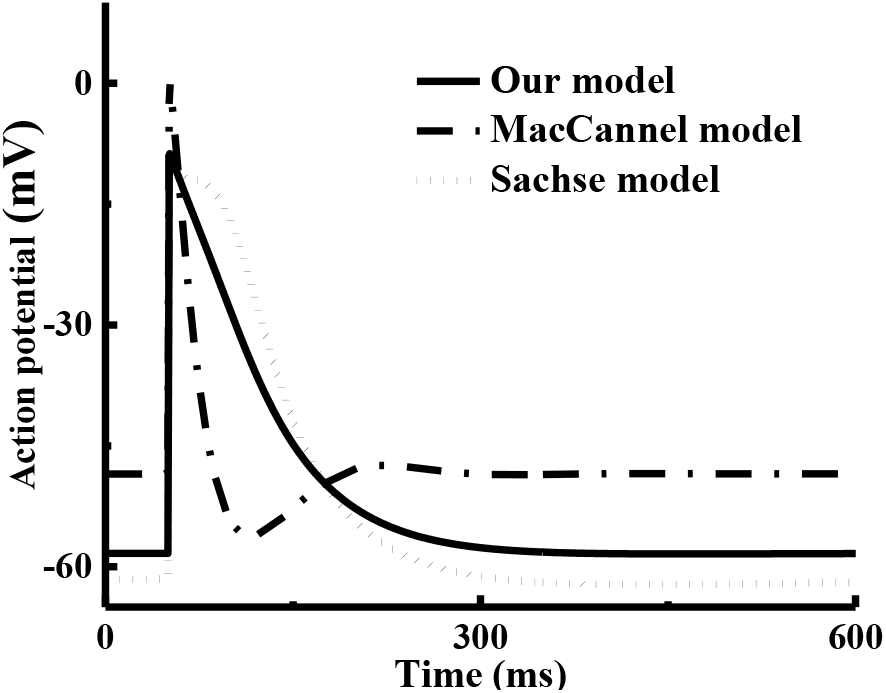
Comparison of action potential among our model, MacCannel model, and Sachse model with a second stimulus current −50pA/pF.

**Table S2.** Compared ionic channel of the different MFB model.

| Ionic channel | MacCannell | Sachse | Novel model | Ref. |
| --- | --- | --- | --- | --- |
| Fast $Na^+$ current | -- | -- | ++ | (1) |
| Mechano-gated current | -- | -- | ++ | (3) |
| Inward rectifying K1 current | ++ | ++ | ++ | (4) |
| Voltage-dependent Kv current | ++ | ++ | ++ | (4) |
| NaK Pump currnet | ++ | -- | ++ | (5) |
| Background current | ++ | ++ | ++ | (4) |

##### Mechanelectrical Model of Left Ventricular CM

Pandit *et al(8)* and ten Tusscher *et al*(9) electrophysiological model of adult rat and human left ventricular CM are widely recognized. We integrated Mullins *et al*(10) mechanic model into the above models to describe the mechanelectrical behavior of human and rat ventricular CM. We did some changes in the integrative mechanelectrical model.

1. The kinetics equation of Ca^2+^ binding to troponin C from Mullins model was added to the Tusscher model to describe excitation-contraction coupling of CM. The model parameters were shown in Table S2.

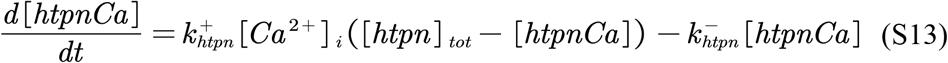

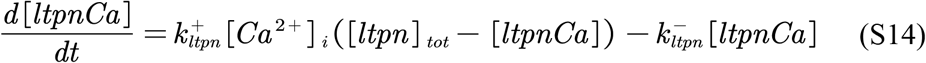

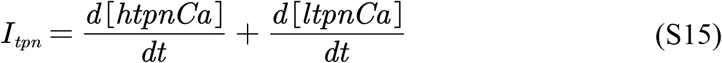

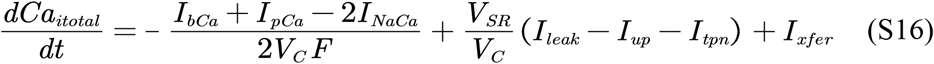
2. The Ca^2+^ transient module of Pandit rat model was replaced by ten Tusscher human model, because the Ca^2+^ transient equations and parameters of Pandit model did not seem to simulate the Ca^2+^ spark (**Fig. S5**). Cytoplasmic buffer Ca^2+^ concentration was revised from 0.2 μM to 0.17 μM in order to agree will with experimental data(11, 12).
3. We corrected the normalized contraction force of Mullins *et al* [6] mechanic model to describe the myofilament contractile in human ventricular CM, based on the newly experimental data. The maximal reference tension was set to 120kPa(13). The simulated results is consistent with similar experiment (14) at long sarcomere length (SL~ 2.25μm) and saturating calcium levels ([Ca^2+^]_o_ = 11uM) (Fig. S6). it’s worth noting that these mechanical experiment parameters are measured at 15°C, because the CM seem to be not enough robust at 37°C(15). According to rat myocardium measurement(16), sarcomere contractile may be higher than recent results at physiological temperature.

**Figure S5.**
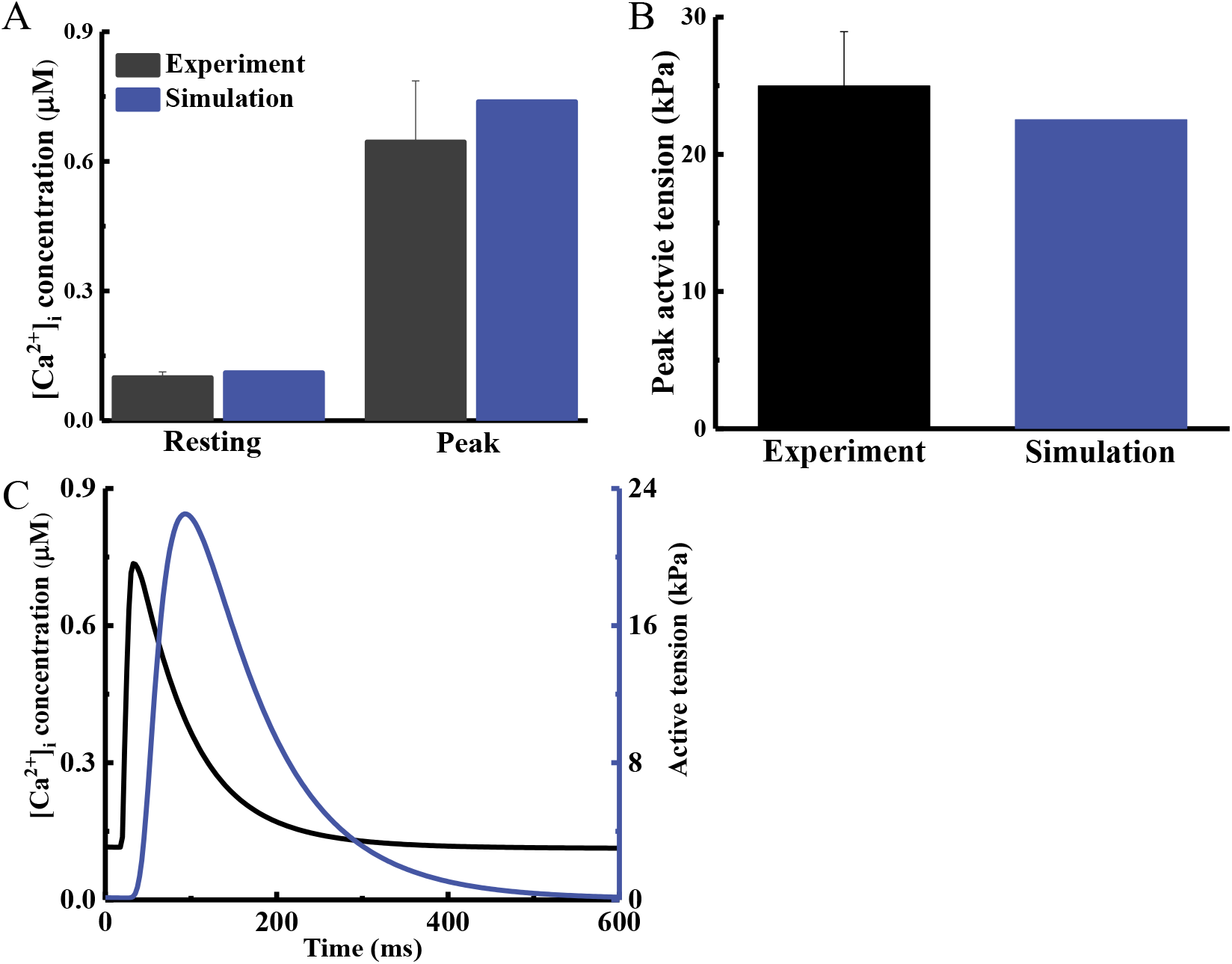
Simulated and experimental resting and peak Ca^2+^ transients of rat CM (A). Active tension of rat CM at initial sarcomere length (SL =1.9 μm) (B). (C) Simulated Ca^2+^ transient and active tension. Experimental data are from Kaprielian *et al* (11) and Kentish *et al*(12).

**Figure S6.**
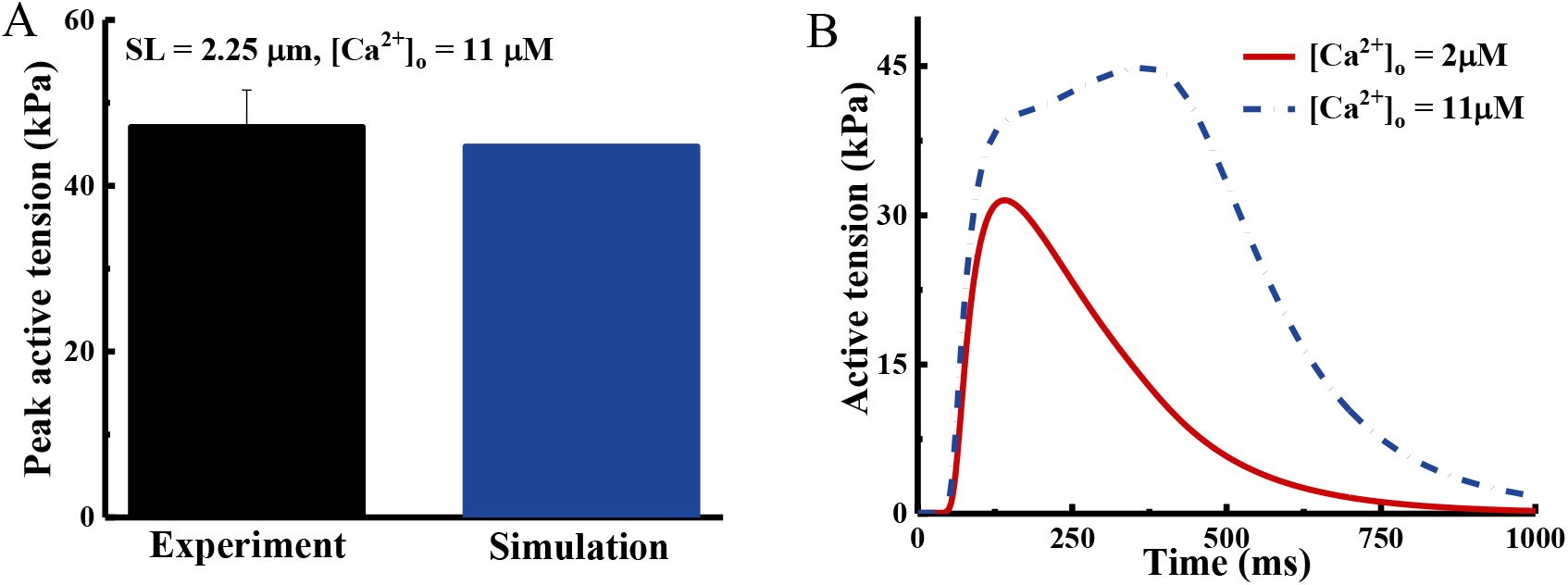
Simulated and experimental peak active tension of human CM at SL = 2.25 μm (A). Active tension of human CM with SL = 2.25 at calcium levels (2 μM *vs*. 11 μM). Experimental data are from Mcdonald *et al* (14).

**Figure S7.**
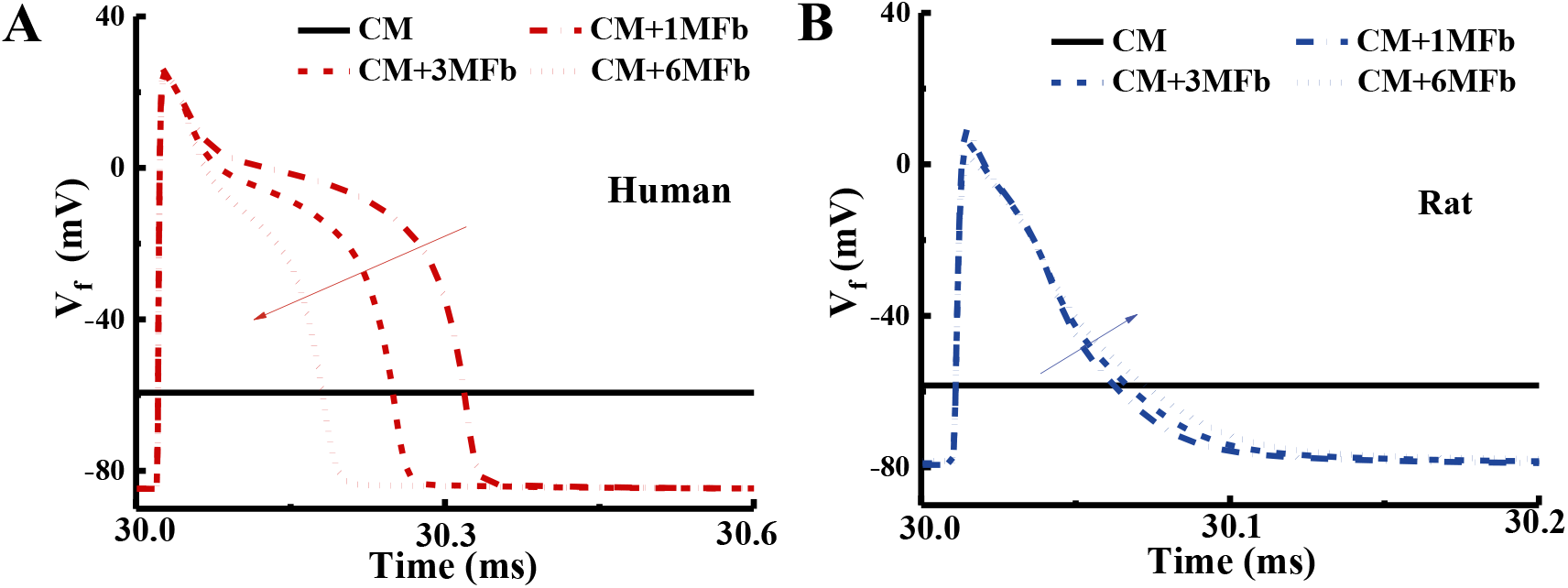
The action potential of MFB coupled human CM (A) and rat CM (B). The trend of action potential of MFB is consistent with the species of CM.

**Figure S8.**
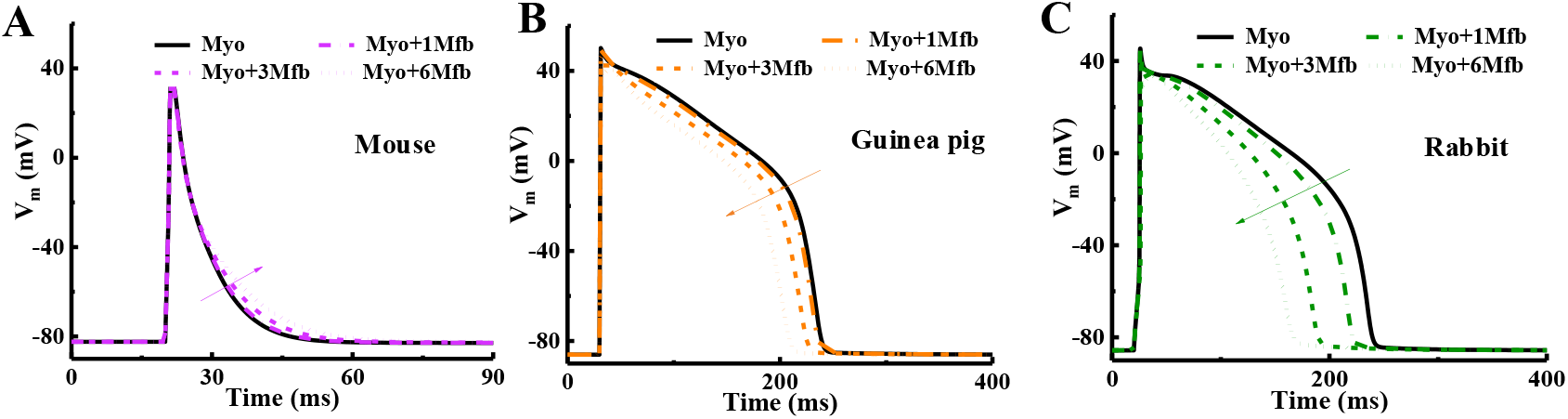
Tends of action potential of mouse (A), guinea pig (B) and rabbit (C) left ventricular CM coupled with zero, one, three or six MFBs. Action potential trends of CM coupled with MFB in rabbit and guinea pig are similar to human and that of mouse is similar to rat.

**Figure S9.**
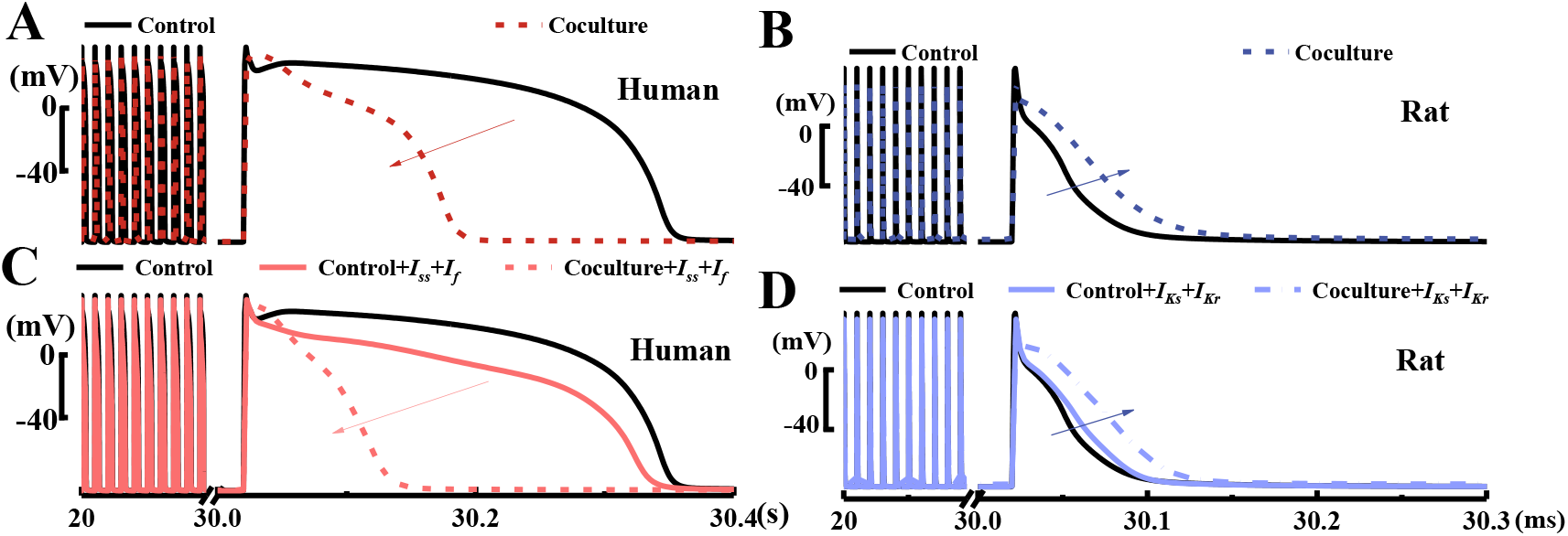
The synergistic coupling shortens the action potential of human CM (A) and prolongs that of rat CM (B). The trends of action potential of human CM and rat CM do not change after replacing rat I_ss_, I_f_ to human I_Kr_, I_Ks_ in human CM (C) and replacing human I_Kr_, I_Ks_ to rat I_ss_, I_f_ in rat CM (D).

**Table S2.**
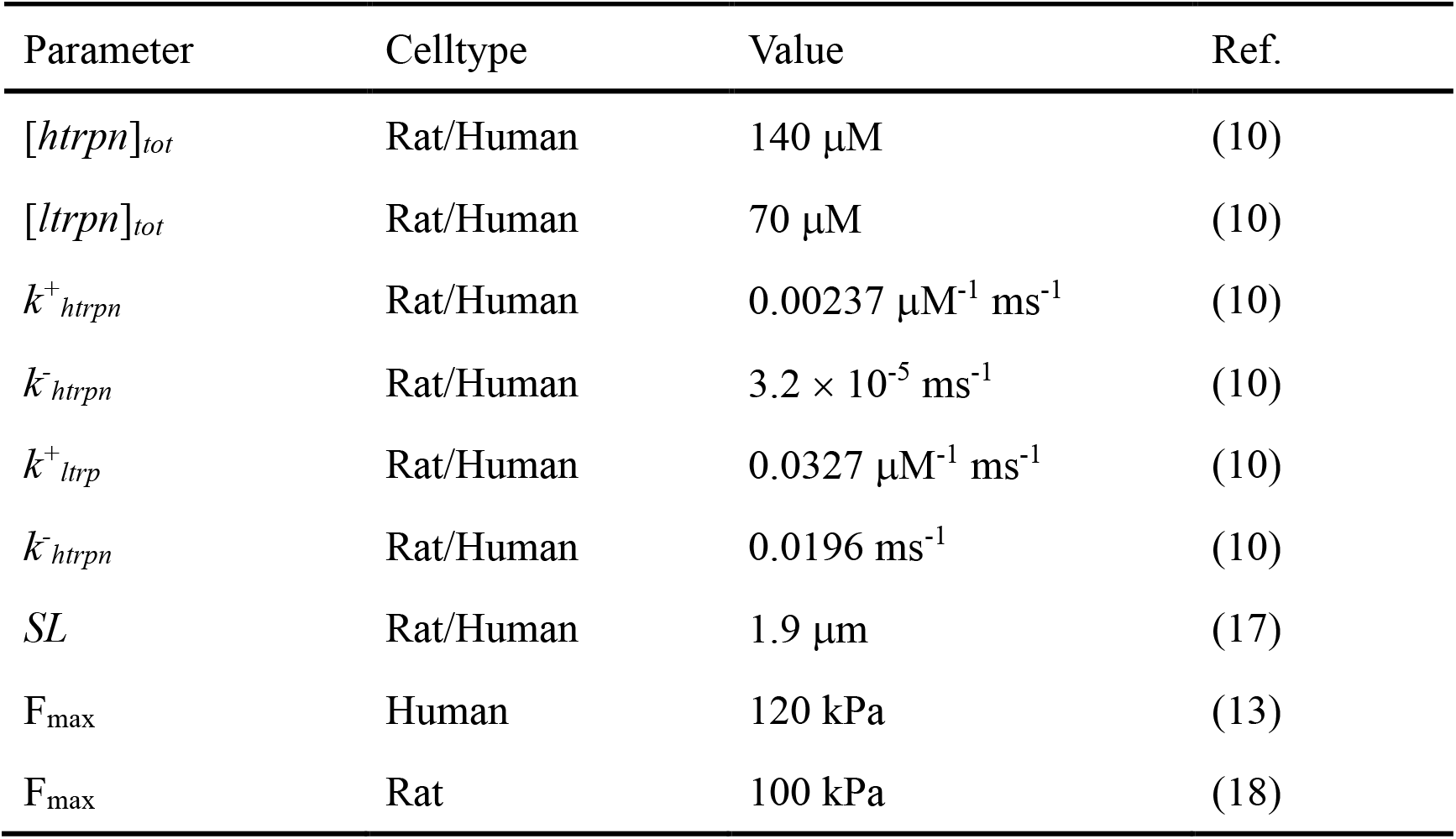
Modified parameters of the mechanelectrical model in human and rat CM

| Parameter | Celltype | Value | Ref. |
| --- | --- | --- | --- |
| $[htrpn]_{tot}$ | Rat/Human | 140 $\mu\text{M}$ | (10) |
| $[ltrpn]_{tot}$ | Rat/Human | 70 $\mu\text{M}$ | (10) |
| $k^+_{htrpn}$ | Rat/Human | 0.00237 $\mu\text{M}^{-1} \text{ms}^{-1}$ | (10) |
| $k^-_{htrpn}$ | Rat/Human | $3.2 \times 10^{-5} \text{ms}^{-1}$ | (10) |
| $k^+_{ltrpn}$ | Rat/Human | 0.0327 $\mu\text{M}^{-1} \text{ms}^{-1}$ | (10) |
| $k^-_{ltrpn}$ | Rat/Human | 0.0196 $\text{ms}^{-1}$ | (10) |
| $SL$ | Rat/Human | 1.9 $\mu\text{m}$ | (17) |
| $F_{\max}$ | Human | 120 kPa | (13) |
| $F_{\max}$ | Rat | 100 kPa | (18) |

